# Pyroptosis-dependent and -independent cross-priming of CD8^+^ T cells by intestinal epithelial cell-derived antigen

**DOI:** 10.1101/2021.07.08.451636

**Authors:** Katherine A. Deets, Randilea D. Nichols, Isabella Rauch, Russell E. Vance

## Abstract

The innate immune system detects pathogens and initiates adaptive immune responses. Inflammasomes are central components of the innate immune system, but whether inflammasomes provide sufficient signals to activate adaptive immunity is unclear. In intestinal epithelial cells (IECs), inflammasomes activate a lytic form of cell death called pyroptosis, leading to epithelial cell expulsion and the release of cytokines. Here we employed a genetic system to show that simultaneous antigen expression and inflammasome activation specifically in IECs is sufficient to activate CD8^+^ T cells. By genetic elimination of direct T cell priming by IECs, we found that IEC-derived antigens are cross-presented to CD8^+^ T cells. However, activation of CD8^+^ T cells by IEC-derived antigen only partially depended on IEC pyroptosis. In the absence of inflammasome activation, cross-priming of CD8^+^ T cells required *Batf3*^+^ dendritic cells (cDC1), whereas cross-priming in the presence of pyroptosis did not. These data suggest the existence of parallel pyroptosis-dependent and pyroptosis-independent but cDC1-dependent pathways for cross-presentation of IEC-derived antigens.

## Introduction

The innate immune system provides a crucial first line of defense against invading pathogens, and in addition, activates and guides subsequent adaptive immune responses. Although the role of innate immunity in promoting adaptive immunity has long been appreciated (Janeway, 1989), most studies have focused on the contributions of Toll-like receptors (TLRs), and significantly less is known about how other innate immune pathways influence adaptive immunity (Iwasaki & Medzhitov, 2015).

Inflammasomes are a heterogeneous group of cytosolic innate immune sensors, each of which oligomerizes in response to specific stimuli, including pathogen-associated molecules and activities or cellular damage (Broz & Dixit, 2016; Rathinam & Fitzgerald, 2016). Regardless of the input signal, a common output of inflammasome activation is the recruitment and activation of caspase proteases (e.g., Caspase-1), which cleave and activate the inflammatory cytokines pro-interleukin (IL)-1β and pro-IL-18 and/or the pore-forming protein gasdermin D. Active gasdermin D oligomerizes in the plasma membrane to form pores that serve as a conduit for the release of active IL-1β and IL-18 and can also initiate pyroptotic cell death and/or lysis (de Vasconcelos, Van Opdenbosch, Van Gorp, Parthoens, & Lamkanfi, 2019; DiPeso, Ji, Vance, & Price, 2017; Evavold et al., 2018; He et al., 2015; Heilig et al., 2018; Kayagaki et al., 2015; Shi et al., 2015). In intestinal epithelial cells (IECs), inflammasome activation also results in the expulsion of cells from the epithelial monolayer into the intestinal lumen. Pyroptosis and cell expulsion provide host defense against intracellular pathogens by eliminating their replicative niche (Fattinger et al., 2021; Hausmann et al., 2020; Mitchell et al., 2020; Rauch et al., 2017; Sellin et al., 2014).

The role of inflammasomes during adaptive immunity remains incompletely understood and inflammasomes appear to have both beneficial and detrimental effects on the adaptive response, depending on the context (Deets & Vance, 2021; Evavold & Kagan, 2018). While there remains limited evidence on how inflammasome activation and pyroptosis impacts presentation of cell-derived antigens, IL-1β and IL-18 have been implicated in driving type 1 helper T cell (Th1), Th17, and CD8^+^ T cell immunity following bacterial infections (Kupz et al., 2012; O’Donnell et al., 2014; Pham et al., 2017; Tourlomousis et al., 2020; Trunk & Oxenius, 2012). Likewise, CD4^+^ and CD8^+^ T cell responses to influenza have been shown to require inflammasome signaling components (Ichinohe, Lee, Ogura, Flavell, & Iwasaki, 2009). However, inflammasome activation has also been found to inhibit T cell-mediated immunity (Sauer et al., 2011; Theisen & Sauer, 2017), including through the pyroptotic destruction of key antigen presenting cells (APCs) (McDaniel, Kottyan, Singh, & Pasare, 2020; Tourlomousis et al., 2020). Inflammasomes have also been suggested to influence adaptive responses to tumors (reviewed in (Deets & Vance, 2021; Evavold & Kagan, 2018)).

To date, most studies have evaluated the effect of inflammasome activation on adaptive immunity in the context of infections. Though they are physiologically relevant, microbial infections are also complex to analyze since they engage multiple innate receptors, including TLRs. It thus remains unknown whether inflammasome activation alone provides sufficient co-stimulatory signals to initiate an adaptive response. In addition, the fate of antigens after inflammasome activation remains poorly understood. Conceivably, expulsion of pyroptotic epithelial cells may result in the loss of antigen, thereby hindering adaptive immunity, or alternatively, pyroptosis may promote the release of antigens to APCs to activate adaptive immunity.

To investigate how inflammasome activation might influence adaptive immunity, we focused on the NAIP–NLRC4 inflammasomes, which specifically respond to flagellin (via NAIP5/6) or bacterial type III secretion system proteins (via NAIP1/2) (Kofoed & Vance, 2011; Rauch et al., 2016; Zhao et al., 2016; Zhao et al., 2011). Although most inflammasomes require the adaptor protein Apoptosis-associated Speck-like protein containing a Caspase-activation and recruitment domain (ASC) to recruit and activate pro-Caspase-1, NLRC4 is able to bind and activate pro-Caspase-1 directly—though ASC has been found to enhance the production of IL-1β and IL-18 (Broz, Newton, et al., 2010; Broz, von Moltke, Jones, Vance, & Monack, 2010; Mariathasan et al., 2004). Cleavage of gasdermin D does not require ASC following NAIP– NLRC4 activation (He et al., 2015; Kayagaki et al., 2015; Shi et al., 2015). NAIPs and NLRC4 are highly expressed in intestinal epithelial cells (IECs), where they provide defense against enteric bacterial pathogens including *Citrobacter* (Nordlander, Pott, & Maloy, 2014), *Salmonella* (Fattinger et al., 2021; Hausmann et al., 2020; Rauch et al., 2017; Sellin et al., 2014) and *Shigella* (Mitchell et al., 2020). Inflammasome-driven IEC expulsion appears to be a major mechanism by which NAIP–NLRC4 provides innate defense against enteric pathogens. However, it is not currently known how pyroptosis and IEC expulsion influence the availability of IEC-derived antigens and what impact this has on the adaptive immune response.

Indeed, even at steady state in the absence of inflammasome activation and pyroptosis, it remains unclear how antigens present in IECs are delivered to APCs to stimulate adaptive immune responses, or whether perhaps IECs can directly activate T cells (Christ & Blumberg, 1997; Nakazawa et al., 2004). Conventional type 1 dendritic cells (cDC1s) are thought to acquire apoptotic bodies from IECs and shuttle the cell-associated antigens through the MHC II pathway to drive a tolerogenic CD4^+^ T cell response under homeostatic conditions (Cummings et al., 2016; Huang et al., 2000). Additionally, in the context of inflammation, a subset of migratory cDC1s have been shown to also take up IEC-derived antigen to activate CD8^+^ T cells; however, it remains unclear how these cDC1s acquire IEC-derived antigen.

Because NAIP–NLRC4 activation can result in IEC pyroptosis prior to the expulsion of IECs from the epithelium (Rauch et al., 2017), we hypothesized that cell lysis could release antigen basolaterally, which could then be taken up by cDC1s and cross-presented to CD8^+^ T cells. To address the role of inflammasome-induced cell death in activation of CD8^+^ T cells, we used a genetic mouse model in which an Ovalbumin (Ova)-Flagellin (Fla) fusion protein is inducibly expressed specifically in intestinal epithelial cells (Nichols, von Moltke, & Vance, 2017). The OvaFla fusion protein provides a model antigenic epitope (SIINFEKL) to activate specific CD8^+^ (OT-I) T cells (Hogquist et al., 1994), concomitant with the activation of the NAIP–NLRC4 inflammasome by a C-terminal fragment of flagellin that does not activate TLR5. This genetic system has the advantage of selectively activating inflammasome responses in the absence of exogenous or pathogen-derived TLR ligands, allowing us to address the sufficiency of inflammasome activation for adaptive responses. Our results suggest the existence of distinct pyroptosis-dependent and pyroptosis-independent pathways for cross-presentation of IEC-derived antigens *in vivo*.

## Results

### Genetic system for NAIP–NLRC4 activation in IECs

We took advantage of a previously established mouse model (Nichols et al., 2017) that allows for Cre-inducible and cell type-specific NAIP–NLRC4 activation (Figure 1A). These mice harbor an *OvaFla* gene fusion that encodes a non-secreted chicken ovalbumin protein—a model antigen—fused to the C-terminal 166 amino acids of flagellin—an agonist of NAIP–NLRC4 but not TLR5 (Nichols et al., 2017). The *OvaFla* gene is inserted within the constitutively expressed *Rosa26* locus, downstream of a floxed transcriptional stop cassette and upstream of an IRES-GFP cassette. To create a genetic system for inducible NAIP–NLRC4 activation in IECs, we crossed the *OvaFla* mice to Villin-Cre-ER^T2^ mice (el Marjou et al., 2004), which harbor a tamoxifen-inducible Cre recombinase driven by the endogenous *Villin* promoter. The resulting *OvaFla* Villin-Cre-ER^T2^ (hereafter shortened to “OvaFla”) mice respond to tamoxifen administration by expressing Cre, and subsequently the OvaFla protein, specifically in IECs. To study the influence of NAIP–NLRC4 activation, pyroptosis, and cytokine production on CD8^+^ T cell activation, we generated *Nlrc4^−/−^*, *Gsdmd^−/−^*, and *Asc*^−/−^ OvaFla lines.

**Figure 1.**
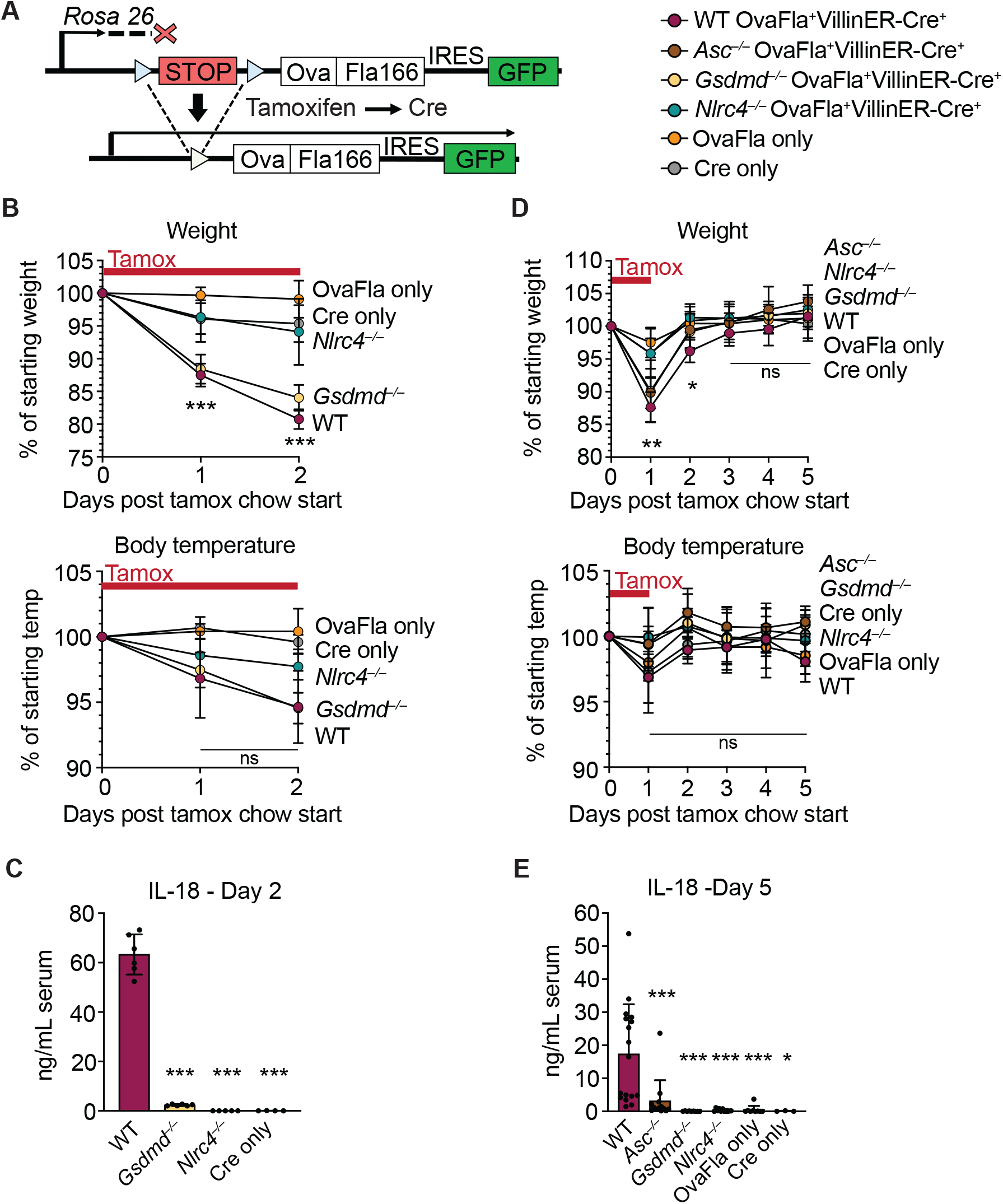
Genetic OvaFla VillinER-CreT2 system results in NAIP–NLRC4 activation in IECs of mice upon tamoxifen chow administration. **A.** Schematic of the OvaFla gene cassette on the Rosa 26 locus. The cassette contains full-length ovalbumin, flagellin with a C-terminal truncation at amino acid 166, and an IRES-GFP. When OvaFla mice are crossed to mice containing the tamoxifen-inducible VillinER-CreT2, tamoxifen administration results in Cre-controlled excision of the stop cassette and expression of the OvaFla fusion protein and GFP within IECs. **B.** Daily weight (top) and rectal temperature (bottom) measurements of OvaFla mice during a two-day course of tamoxifen chow (depicted as red bar). **C.** Quantification of IL-18 ELISA performed on serum from the mice shown in panel B at day 2 post tamoxifen chow start. Each dot represents an individual mouse. **D.** Daily weight (top) and rectal temperature (bottom) measurements of OvaFla mice following a single day pulse of tamoxifen chow (depicted as red bar). **E.** Quantification of IL-18 ELISA performed on serum from the mice shown in panel D at day 5 post tamoxifen chow start. Each dot represents an individual mouse. B-E, data shown as mean +/– SD. Significance calculated using one-way ANOVA and Tukey’s multiple comparisons test (\**p* < 0.05, \*\**p* <0.01, \*\*\**p* < 0.001). D, B, *p* valutes between WT and *Nlrc4*^−/−^ are shown. C, E, *p* values between WT and other experimental groups are shown.

Tamoxifen is typically administered in a corn oil emulsion through oral gavage or intraperitoneal injection. In preliminary experiments, we found corn oil contains trace bacterial contaminants that activate TLR signaling (Nichols, 2017). Thus, to avoid confounding effects of TLR activation, and to isolate the specific effects of inflammasome activation, we administered tamoxifen orally through a commercially available tamoxifen-containing chow. OvaFla mice were fed *ab libitum*, and their weight and temperature were tracked daily as previously-described indicators of NAIP–NLRC4 activation (von Moltke et al., 2012). After a single day on the tamoxifen diet, WT OvaFla and *Gsdmd*^−/−^ OvaFla mice lost a significant amount of weight, and by day two of the tamoxifen diet, these mice needed to be euthanized due to exceeding the humane weight loss endpoint on our animal protocol (Figure 1B, top). In contrast, the *Nlrc4*^−/−^ OvaFla mice, as well as the OvaFla-only and Cre-only littermate control mice, maintained a consistent body weight and appeared healthy over the two-day time course. Although not statistically significant, the WT OvaFla and *Gsdmd*^−/−^ OvaFla mice also exhibited decreases in core body temperature by day two relative to the *Nlrc4*^−/−^ OvaFla mice (Figure 1B, bottom), consistent with previous analyses using recombinant flagellin protein (FlaTox) to induce acute NAIP–NLRC4 activation (Rauch et al., 2017; von Moltke et al., 2012).

Serum was collected from OvaFla mice at day two of the tamoxifen diet and assayed for IL-18, which is released from IECs following NAIP–NLRC4 activation (Rauch et al., 2017). The serum of WT OvaFla mice contained approximately 60 times more IL-18 than the serum of *Gsdmd*^−/−^ OvaFla mice, demonstrating that gasdermin D is required for IL-18 release from IECs following NAIP–NLRC4 activation (Figure 1C). IL-18 was not detected in the *Nlrc4*^−/−^ mice or in the OvaFla-only or Cre-only littermate controls. Taken together, these data show that the OvaFla system results in robust NAIP–NLRC4 activation in IECs following tamoxifen administration.

To study the CD8^+^ T cell response to IEC-derived antigen following NAIP–NLRC4 activation, we shortened the administration of tamoxifen chow to a single day pulse to limit confounding effects of morbidity in the NAIP–NLRC4 sufficient strains. We again monitored weight and rectal temperature each day. We found that while the WT, *Asc^−/−^*, and *Gsdmd*^−/−^ OvaFla mice initially lost weight, weight loss was reversed by day three post tamoxifen chow start (Figure 1D, top). No significant difference in core body temperature was found between strains over the five-day experiment (Figure 1D, bottom).

Serum was collected at day five post start of the tamoxifen chow diet and again assayed for IL-18 through ELISA. Similar to the two-day tamoxifen pulse, a single day of tamoxifen chow resulted in significant IL-18 production in the WT OvaFla mice but minimal to no detectable IL-18 in the other OvaFla strains (Figure 1E). The WT mice exhibited heterogeneity in the IL-18 response with the single day chow pulse, which may be related to some mice being averse to consuming the tamoxifen chow (Chiang et al., 2010) or heterogeneity in the kinetics of the response.

We also performed immunofluorescence imaging of the small intestines of mice from each of the OvaFla lines at day two following the start of a single day pulse of tamoxifen chow. The presence of an IRES-GFP downstream of the *OvaFla* gene allows us to track the expression of the transgene. While approximately 30% of the IECs were GFP^+^ in *Nlrc4*^−/−^ OvaFla mice, only about 2% of the IECs were GFP^+^ in the WT, *Asc^−/−^*, or *Gsdmd*^−/−^ OvaFla mice at that time point (Figure 2). This result was anticipated because previous work (Rauch et al., 2017; Sellin et al., 2014) found that IECs are rapidly expelled from the epithelium upon NAIP–NLRC4 activation. Given that we observe robust IL-18 levels in the serum of WT mice (Figure 1C, E), we believe the transgene is expressed in WT mice, but NLRC4^+^ cells that express high levels of the transgene will be expelled, limiting our ability to detect them. Although pyroptosis of IECs requires Gasdermin D, NAIP–NLRC4-induced IEC expulsion was previously found to be independent of Gasdermin D, likely due to the existence of an NLRC4-Caspase-8-dependent apoptosis pathway that also leads to IEC expulsion (Man et al., 2013; Rauch et al., 2017). These previous results are consistent with the apparent loss of GFP^+^ cells we observe in the *Gsdmd*^−/−^ OvaFla mice.

**Figure 2.**
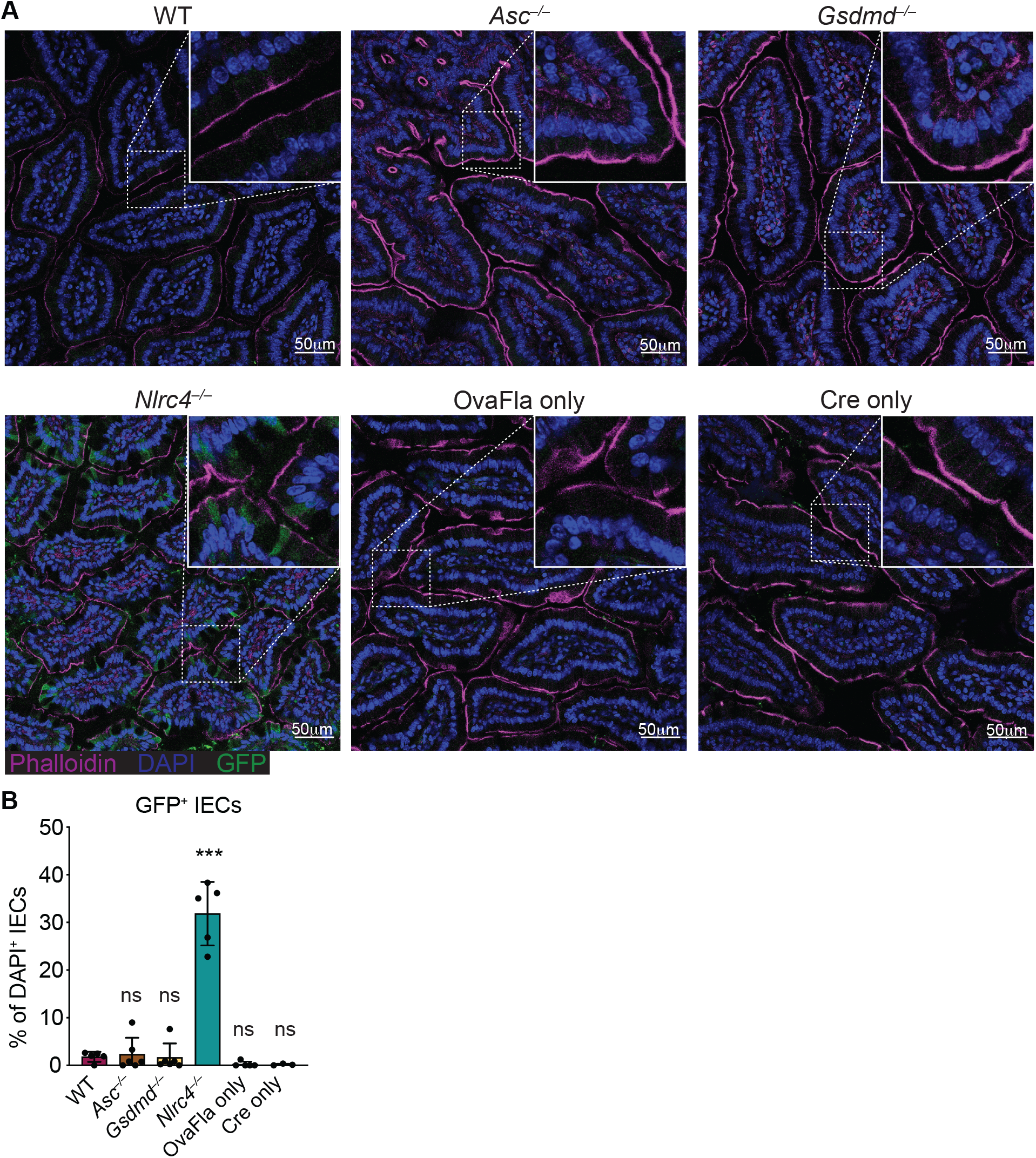
GFP^+^ cells accumulate in *Nlrc4^−/−^* mice following tamoxifen administration. **A.** Representative immunofluorescence images of the small intestines of indicated OvaFla mice on day 2 following a single day pulse of tamoxifen chow. **B.** Quantification of DAPI+ IECs that are also GFP+ for each OvaFla line. Data are from two independent experiments, and each dot represents an individual mouse. Data shown as mean+/– SD. Significance calculated using one-way ANOVA and Tukey’s multiple comparisons test (\**p* < 0.05, \*\**p* < 0.01, \*\*\**p* < 0.001). Only *p* values between WT and other experimental groups are shown.

Taken together, these data show that OvaFla production under control of the tamoxifen-inducible Villin-Cre-ER^T2^ system results in robust NAIP–NLRC4 activation in the IECs of mice. A single day pulse of tamoxifen chow leads to significant IL-18 production without gross morbidity or mortality in the NAIP–NLRC4 sufficient strains. Additionally, OvaFla likely accumulates in the IECs of the *Nlrc4*^−/−^ OvaFla mice, as these cells do not undergo NAIP– NLRC4-driven cell expulsion.

### CD8^+^ T cell activation by epithelial antigens

We followed the response of Ova-specific TCR transgenic OT-I CD8^+^ T cells following OvaFla induction in each of our mouse lines. Congenically marked (CD45.1^+^ or CD45.1^+^ CD45.2^+^) OT-I T cells were harvested from the spleens and mesenteric lymph nodes of OT-I *Rag2*^−/−^ mice, labeled with CellTrace Violet proliferation dye, and intravenously transferred into the OvaFla mice (2×10^4^ cells per mouse) (Figure 3A). Immediately following adoptive transfer, the mice were placed on tamoxifen chow for a single day. At day five post adoptive transfer, the mice were euthanized, and their mesenteric lymph nodes, which drain immune cells from the intestines (Esterhazy et al., 2019), and spleens were analyzed for OT-I T cell proliferation and activation.

**Figure 3.**
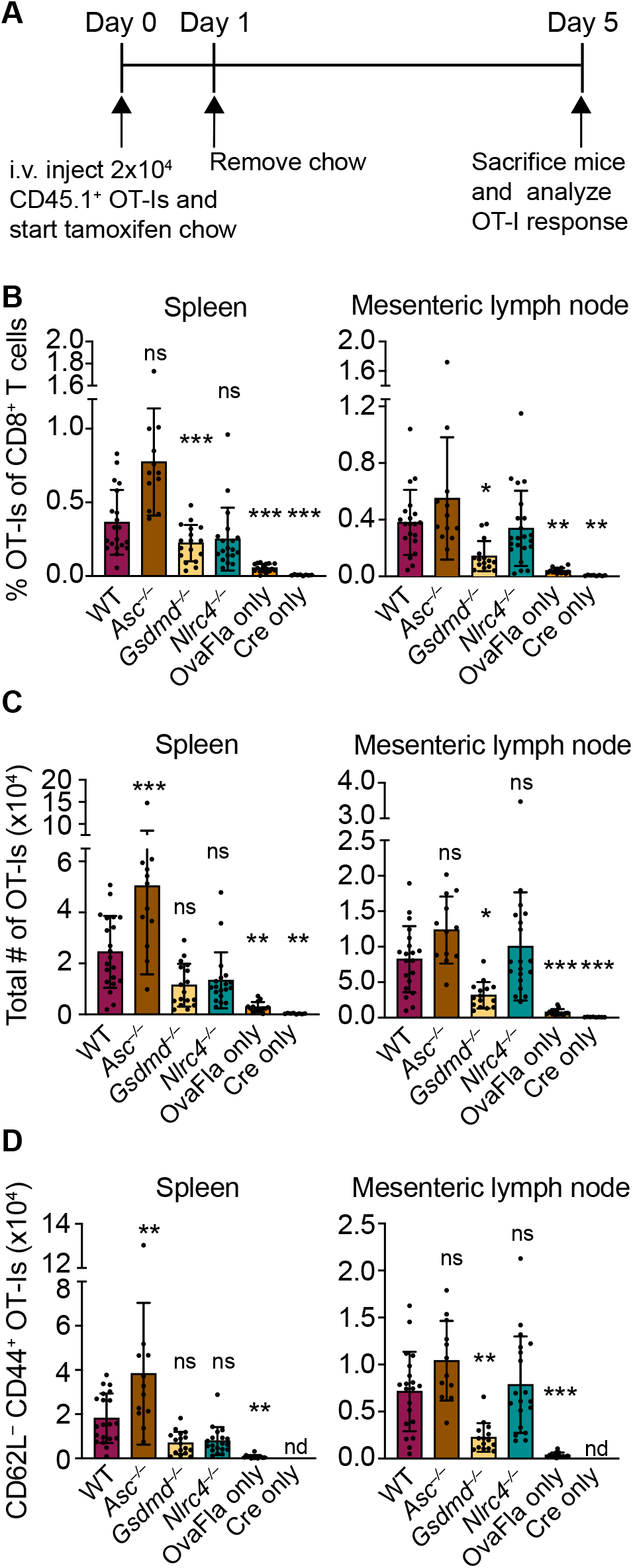
OvaFla expression in IECs results in OT-I proliferation and activation that is independent of ASC and NLRC4 but partially dependent on gasdermin D. **A.** Overview of experimental setup for analyzing OT-I responses to OvaFla production in IECs. **B.** Quantification of OT-Is as a percent of total CD8+ T cells per spleen (left) and mesenteric lymph node (right) **C.** Total number of OT-Is per spleen (left) and mesenteric lymph node (right). **D.** Total number of CD64L^−^CD44^+^ OT-Is per spleen (left) and mesenteric lymph node (right). B-D, data are from three independent experiments, and each dot represents an individual mouse. Data shown as mean +/– SD. Significance calculated using one-way ANOVA and Tukey’s multiple comparisons test (\**p* < 0.05, \*\**p* < 0.01, \*\*\**p* < 0.001). Only *p* values between WT and other experimental groups are shown.

A dividing OT-I population was identified by flow cytometry in each Cre^+^ OvaFla line (Figure 3–figure supplement 1A, 1B). These data demonstrate that antigens expressed in IECs can be processed and presented to activate CD8^+^ T cells *in vivo*. Surprisingly, however, there was minimal difference in the relative percent (Figure 3B), absolute number (Figure 3C), or activation status (defined as CD62L^−^CD44^+^)(Figure 3D, Figure 3–figure supplement 1C, D) of OT-I T cells between the WT and *Nlrc4*^−/−^ OvaFla mice in either the spleen or mesenteric lymph node. In fact, relative to the WT OvaFla mice, a higher percent of the OT-I T cells in the *Nlrc4*^−/−^ OvaFla mice produced IFNγ and TNFα following *ex vivo* stimulation with PMA and ionomycin (Figure 3–figure supplement 1E). These data indicate OT-I T cells respond to IEC-expressed Ova in a manner that is independent of NAIP–NLRC4 activation. However, the specific lack of IEC expulsion and the resulting higher accumulation of antigen in IECs in *Nlrc4*^−/−^ mice (Figure 2) means that the WT and *Nlrc4*^−/−^ mice are not truly comparable.

In contrast to *Nlrc4*^−/−^ IECs, both *Asc*^−/−^ and *Gsdmd*^−/−^ IECs are expelled after inflammasome activation and thus exhibit similarly low OvaFla-IRES-GFP transgene expression in IECs as seen in WT mice (Figure 2). Both strains are also defective for IL-18 release (Figures 1C, 1E). The major difference between the two strains is that *Asc*^−/−^ cells can still undergo *Gsdmd*-dependent pyroptosis, whereas *Gsdmd*^−/−^ cells do not undergo lytic pyroptosis but are nevertheless expelled from the epithelium as intact apoptotic cells, likely via a Caspase-1 and/or −8 pathway (Man et al., 2013; Rauch et al., 2017). There was little difference in OT-I numbers (Figures 3B, 3C) or activation (Figure 3D, Figure 3–figure supplement 1C, D), as well as no difference in IFNγ and TNFα production (Figure 3–figure supplement 1E), between the WT OvaFla mice and the *Asc*^−/−^ OvaFla mice. However, we were surprised to observe partial deficits in OT-I responses in *Gsdmd*^−/−^ OvaFla mice relative to the other mouse lines (Figures 3B–D, Figure 3–figure supplement 1D), though the deficits were not statistically significant for all parameters measured. Thus, these results suggest inflammasome activation in IECs is not essential for OT-I CD8^+^ T cell activation, yet Gasdermin D-mediated pyroptosis of IECs may play a partial role (see Discussion).

### Cross-presentation of IEC antigens

IECs express MHC class I on their surface and are capable of directly presenting antigen to CD8^+^ T cells (Christ & Blumberg, 1997; Nakazawa et al., 2004). It is therefore possible that the OT-I activation seen in the OvaFla mice is a result of direct presentation of Ova peptide by the IECs expressing OvaFla. However, it is also possible that the OT-I T cells are being cross-primed by cDC1s that engulf and “cross present” the IEC-derived Ova (Cerovic et al., 2015; Z. Liu & Lefrancois, 2004). The fate of IEC-derived antigens and the role of antigen-presentation pathways leading to CD8^+^ T cell activation has not previously been addressed with a completely *in vivo* system that can genetically distinguish cross from direct presentation of IEC antigens.

To determine whether the OT-Is are being activated through cross presentation or direct presentation of Ova peptide, we took advantage of the H-2K^bm1^ mouse model that contains a seven base pair mutation in the gene encoding K^b^ (Schulze et al., 1983). The bm1 mutation renders K^b^ unable to bind the Ova-derived OT-I agonist peptide, SIINFEKL (Nikolic-Zugic & Bevan, 1990). We bred H-2K^bm1^ mice to each of our OvaFla lines to establish mice that make OvaFla in their IECs but are incapable of directly presenting the SIINFEKL peptide (H-2K^bm1+^ OvaFla mice, referred to here as bm1^+^ OvaFla mice). We then generated bone marrow chimeras using bm1^+^ OvaFla mice as lethally irradiated recipients that were reconstituted with WT H-2K^b^ bone marrow from B6 CD45.1 donors (Figure 4A, left). In these chimeras, the IECs produce OvaFla following tamoxifen administration, but the IECs themselves are unable to present SIINFEKL to OT-I T cells. The donor-derived hematopoietic cells, including cross-presenting cDC1s, do not contain the *OvaFla* gene cassette but are able to cross-present the SIINFEKL peptide if they acquire it from IECs (Figure 4A, right). Therefore, in the bm1^+^ OvaFla chimeras, we will only see OT-I proliferation and activation if the SIINFEKL peptide is cross-presented.

**Figure 4.**
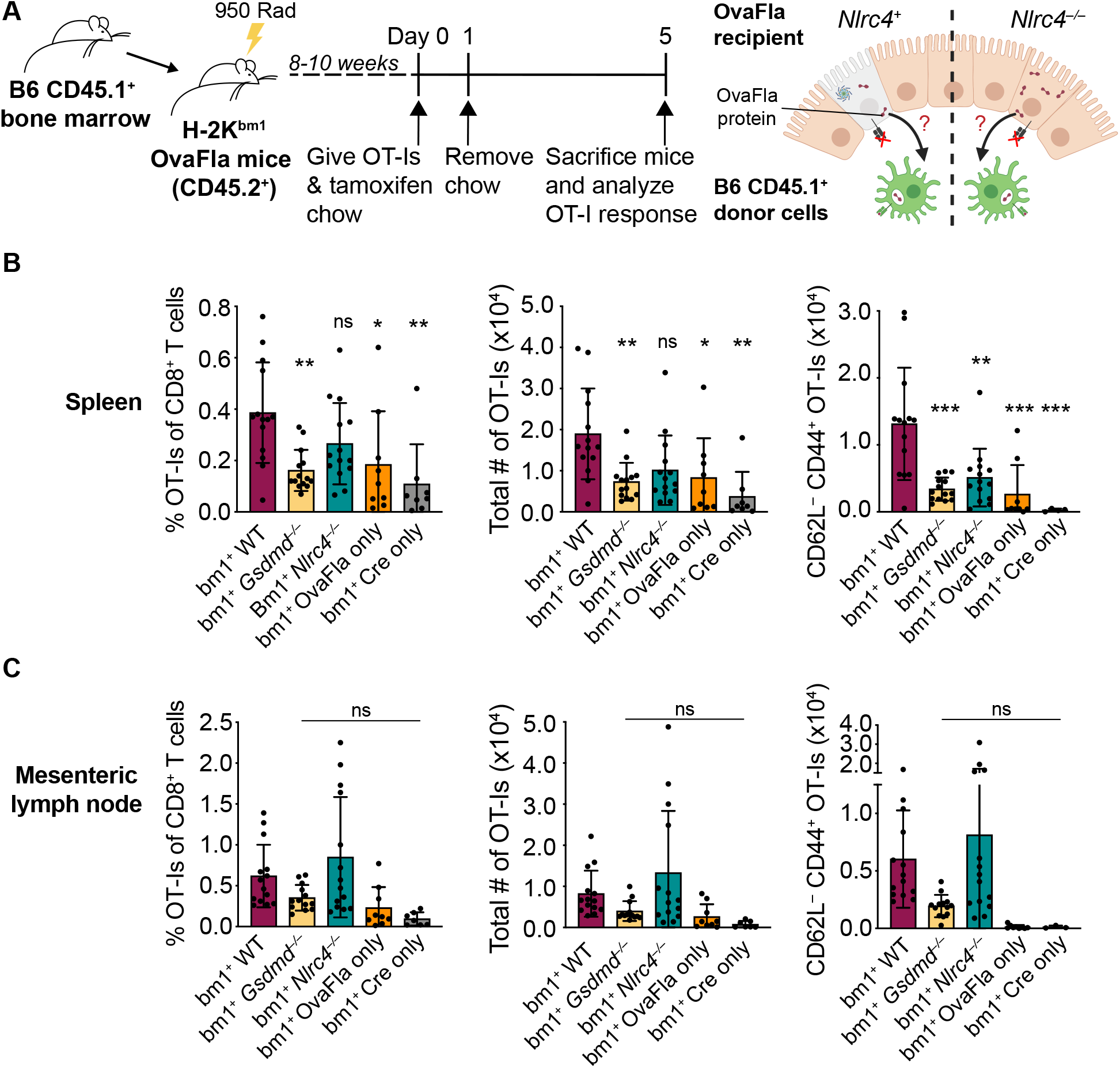
OvaFla expression in IECs results in OT-I cross-priming that is independent of NLRC4 but partially dependent on gasdermin D. **A.** Schematic depicting the production and analysis workflow of chimeric bm1^+^ OvaFla mice (left). At the right, an illustration of either bm1^+^ WT OvaFla mice (left of the dashed line) or bm1^+^ *Nlrc4*^−/−^OvaFla mice (right of the dashed line) following lethal irradiation and reconstitution with bone marrow from B6.SJL mice. **B.** Quantification of OT-Is as a percent of total CD8^+^ T cells (left), the total number of OT-Is (middle), and the total number of CD64L^−^CD44^+^ OT-Is (right) in the spleen. **C.** Quantification of OT-Is as a percent of total CD8^+^ T cells (left), the total number of OT-Is (middle), and the total number of CD64L^−^CD44^+^ OT-Is (right) in the mesenteric lymph nodes. B-C, data are from three independent experiments, and each dot represents an individual mouse. Data shown as mean +/– SD. Significance calculated using one-way ANOVA and Tukey’s multiple comparisons test (\**p* < 0.05, \*\**p* < 0.01, \*\*\**p* < 0.001). Only *p* values between WT and other experimental groups are shown.

Eight to ten weeks after lethal irradiation and reconstitution, bm1^+^ OvaFla mice received 2×10^4^ CD45.1^+^ CD45.2^+^ CellTrace Violet labeled OT-I T cells intravenously and were given a one-day pulse of tamoxifen chow (Figure 4A, left). The mice were euthanized at day five post OT-I transfer, and their spleens and mesenteric lymph nodes were analyzed for OT-I proliferation and activation. Serum was also collected for IL-18 ELISA to confirm NAIP– NLRC4-dependent IL-18 release following OvaFla induction (Figure 4–figure supplement 1A).

Like the non-chimera experiments, an obvious, dividing and activated OT-I population was observed by flow cytometry in each of the OvaFla mouse lines (Figures 4B, C, Figure 4– figure supplement 1B, C). This population was absent in mice given H-2K^bm1^ bone marrow (Figure 4–figure supplement 2), confirming the requirement for APCs to express K^b^ to activate OT-I T cells. These data provide formal genetic evidence that IEC-derived antigens can be cross-presented to activate CD8^+^ T cells *in vivo*.

In the spleen, the bm1^+^ WT and bm1^+^ *Nlrc4*^−/−^ OvaFla mice harbored significantly more OT-I T cells than the bm1^+^ *Gsdmd*^−/−^ OvaFla mice, or the bm1^+^ OvaFla-only and bm1^+^ Cre-only littermate controls, by both percent (Figure 4B, left) and total number (Figure 4B, middle). There were also significantly more activated (CD62L^−^CD44^+^) OT-I T cells in the spleens of bm1^+^ WT mice as compared to bm1^+^ *Gsdmd*^−/−^ mice (Figure 4B, left, Figure 4–figure supplement 1C). In the mesenteric lymph nodes, the relative percent (Figure 4C, left) total number (Figure 4C, middle), and number of activated (Figure 4C, right, Figure 4–figure supplement 1C) OT-I T cells in the bm1^+^ WT OvaFla mice trended higher than the bm1^+^ *Gsdmd*^−/−^ mice and the controls, but was not statistically significant. The reason for the weak responses in the mesenteric lymph nodes is unclear, but others have previously noted negative impacts in irradiation chimeras on the expansion of adoptively transferred OT-I T cells (Kurts, Kosaka, Carbone, Miller, & Heath, 1997). There was no difference across the genotypes in the percent of cells that produce IFNγ and TNFα following *ex vivo* stimulation with PMA and ionomycin (Supp fig 2D).

Taken together, these data provide genetic evidence that OT-I T cells are cross-primed from IEC-derived antigen following OvaFla induction. This cross-priming does not strictly require NAIP–NLRC4 activation but Gasdermin D-induced pyroptosis can promote CD8+ T cell responses, at least for splenic OT-I T cells.

### Pyroptotic cell antigens are cross-presented independent of *Batf3^+^* cDC1s

*Batf3* is a transcription factor required for development of cDC1s (Edelson et al., 2010), and previous work showed that *ex vivo* isolated cDC1s can cross-prime CD8^+^ T cells with IEC-derived antigen (Cerovic et al., 2015). To investigate whether the cross-priming of IEC-derived antigen occurring in the OvaFla mice requires cDC1 cells, we took advantage of our H-2K^bm1^ bone marrow chimera system and compared bm1^+^ OvaFla recipients that received either B6 CD45.1 bone marrow or bone marrow from *Batf3*^−/−^ mice.

As with the above experiments, bone marrow chimeras were made by lethally irradiating bm1^+^ OvaFla mice and transferring donor bone marrow from either B6 CD45.1 or *Batf3*^−/−^ donors. Eight to ten weeks post irradiation, 2×10^4^ CD45.1^+^ CD45.2^+^ CellTrace Violet labeled OT-I T cells were adoptively transferred intravenously, and the mice were given a one-day pulse of tamoxifen chow (as in Figure 4A, left). The mice were sacrificed five days later, and their spleens and mesenteric lymph nodes were analyzed for OT-I proliferation and activation. We confirmed an absence of cDC1 cells in the OvaFla mice that received *Batf3*^−/−^ donor bone marrow (Figure 5A, Figure 5–figure supplement 1).

**Figure 5.**
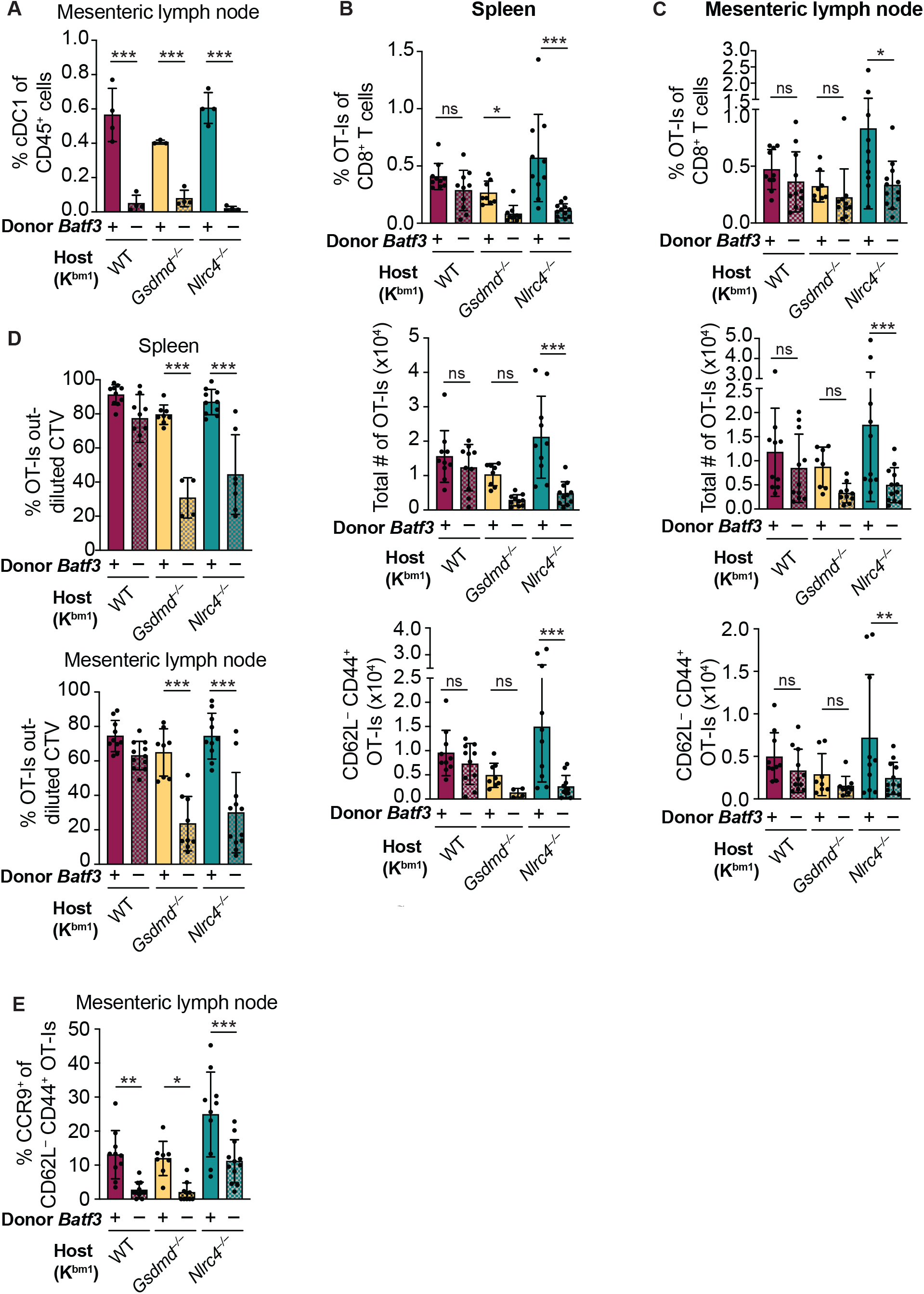
Cross-priming of OT-Is is independent of *Batf3^+^* cDC1s following NAIP–NLRC4 activation in IECs. **A.** Percent of CD45+ cells that are cDC1s in bm1 chimera mice that received either *Batf3^+^* or *Batf3^−^* donor bone marrow. **B.** Quantification of OT-Is as a percent of total CD8^+^ T cells (left), the total number of OT-Is (middle), and the total number of CD64L^−^CD44^+^ OT-Is (right) in the spleen. **C.** Quantification of OT-Is as a percent of total CD8^+^ T cells (left), the total number of OT-Is (middle), and the total number of CD64L^−^CD44^+^ OT-Is (right) in the mesenteric lymph nodes. **D.** Quantification of OT-Is that have out-diluted the CellTrace Violet dye in the spleen (top) and mesenteric lymph nodes (bottom). **E.** Percent of CD64L^−^CD44^+^ OT-Is in the mesenteric lymph node that are CCR9^+^. A, data are from a single experiment. B-D, data are from three independent experiments. E, data are from two independent experiments. Each dot represents an individual mouse. Data shown as mean ¬ ± SD. Significance calculated using one-way ANOVA and Šídák’s multiple comparisons test (\**p* < 0.05, ** *p* <0.01, \*\*\**p* < 0.001).

To our surprise, there was no difference in the relative (Figures 5B–C, top) or total (Figures 5B–C, middle) number of OT-I T cells between bm1^+^ WT OvaFla mice that received B6 CD45.1 or *Batf3*^−/−^ bone marrow in either the spleen or mesenteric lymph node. OT-I T cells in these two mouse groups also appeared to proliferate similarly (Figure 5D, Figure 5–figure supplement 2). Additionally, there was no difference in the percent (Figure 5–figure supplement 3) or total number (Figures 5B–5C, bottom) of CD44^+^ CD62L^−^ OT-I T cells between these two groups. These data suggest a *Batf3*-independent population of DCs are responsible for cross-presentation of IEC-derived antigen following NAIP–NLRC4 activation.

The above findings with WT OvaFla mice are in stark contrast to the *Nlrc4*^−/−^ OvaFla mice, which exhibit a significant decrease in the relative (Figures 5B–C, top) and total (Figures 5B–C, middle) number of OT-I T cells in the spleens and mesenteric lymph nodes of mice that received *Batf3*^−/−^ donor cells compared to the mice that received B6 CD45.1 donor cells. Correspondingly, there was a significant decrease in the total number of CD44^+^ CD62L^−^ OT-I T cells (Figures 5B–C, bottom). The difference in OT-I numbers between these two groups of mice may be related to a relative decrease in proliferation of the OT-I T cells in the mice receiving *Batf3*^−/−^ bone marrow, as evidenced by less dilution of the CellTrace Violet dye (Figure 5D, Figure 5–figure supplement 2).Taken together these data indicate that in the absence of NLRC4-inflammasome activation, efficient cross-presentation of IEC-derived antigen *in vivo* requires cDC1s, but that this requirement is circumvented in the presence of NLRC4 activation.

NLRC4 activation might promote alternative (cDC1-independent) cross-presentation pathways by the pyroptotic release of antigen and/or inflammatory cytokines. To test whether Gasdermin D is required for cDC1-independent cross-priming, we examined bm1^+^*Gsdmd*^−/−^ chimeras reconstituted with *Batf3*^+^ or *Batf3*^−/−^ bone marrow. The bm1^+^ *Gsdmd*^−/−^ OvaFla mice exhibit a phenotype that falls between the bm1^+^ WT and bm1^+^ *Nlrc4*^−/−^ OvaFla mice, with the only significant differences between WT and *Batf3*^−/−^ bone marrow recipients in the division of OT-I T cells (Figure 5D, Figure 5–figure supplement 2) and relative percent of OT-I T cells in the spleen (Figure 5B, top). These data suggest that cross-priming of OT-I T cells is largely dependent on *Batf3*^+^ cDC1s in the absence of NAIP–NLRC4 activation, but NAIP–NLRC4 activation reveals a *Batf3*-independent cross-presentation pathway.

Regardless of the bone marrow donor, OT-I T cells in the bm1^+^ WT, bm1^+^ *Gsdmd^−/−^*, and bm1^+^ *Nlrc4*^−/−^ OvaFla mice all showed similar levels of TNFα and IFNγ production following *ex vivo* stimulation with PMA and ionomycin (Figure 5–figure supplement 3B). However, when we looked at CCR9 expression as a readout of whether the OT-I T cells were homing to the intestine (Svensson et al., 2002), we found a significant decrease in the number of cells expressing CCR9 in the *Batf3*^−/−^ recipients relative to the B6 CD45.1 recipients across all three mouse lines (Figure 5E). In summary, our data indicate the existence of two potential pathways by which IEC-derived antigens are cross-presented to CD8^+^ T cells: one that requires *Batf3*^+^ cDC1s, and one that does not. However, the *Batf3*^+^ cDC1s appear necessary for instructing antigen specific CD8^+^ T cells back to the intestine.

## Discussion

Intestinal epithelial cells (IECs) represent an important barrier surface that protects against enteric pathogens. At the same time, IECs also represent a potential replicative niche for pathogens. As such, the immune system must survey IECs for foreign antigens and present those antigens to activate protective adaptive immune responses. In general, it remains poorly understood whether and how IEC-derived antigens are presented to activate T cell responses. In particular, the relative contributions of direct versus cross-presentation of IEC antigens to CD8^+^ T cells has not been thoroughly investigated. Here we employed a genetic system that inducibly expresses a model antigen (ovalbumin) fused to a NAIP–NLRC4 agonist (flagellin) within the cytosol of cells (Nichols et al., 2017). We crossed these “OvaFla” mice to Villin-Cre-ER^T2^ mice, allowing for tamoxifen-inducible expression specifically in IECs. By additionally crossing to an H-2K^bm1^ background (Nikolic-Zugic & Bevan, 1990; Schulze et al., 1983), and using the resulting mice as irradiated recipients for wild-type K^b^ hematopoietic donor cells, we engineered a system in which an IEC-derived ovalbumin antigen (SIINFEKL) cannot be directly presented to OT-I T cells but can still be acquired by hematopoietic cells and cross-presented. Using this system, we established *in vivo* that there is an antigen-presentation pathway in which IEC-derived antigens are cross-presented to activate CD8^+^ T cells. This finding extends previous work indicating that *ex vivo* isolated DCs can cross present IEC-derived antigens to CD8^+^ T cells (Cerovic et al., 2015; Cummings et al., 2016). We show these antigens can activate antigen-specific CD8^+^ T cells *in vivo*, and that this activation can occur even when direct presentation is genetically eliminated. We suggest that the cross-presentation pathway revealed by our analyses could be of importance during infection with pathogens that replicate in IECs, though future studies will be required to evaluate this.

Our genetic system also allowed us to assess the contribution of IEC inflammasome activation to the adaptive immune response. Inflammasomes are a critical component of the innate immune response to many pathogens, and their activation is known to influence adaptive immunity (Deets & Vance, 2021). However, in previous studies, it has been difficult to isolate the specific effects of inflammasome activation on adaptive immunity because microbial pathogens activate numerous innate immune signaling pathways over the course of an infection. By providing a genetically encoded antigen and inflammasome stimulus, we were able to overcome this issue and specifically address the role of inflammasomes in adaptive CD8^+^ T cell responses *in vivo*. We crossed our OvaFla Villin-Cre-ER^T2^ mice to mice deficient in key inflammasome components. Consistent with previous work, we found that *Nlrc4*^−/−^ mice entirely lack the inflammasome response to cytosolic flagellin, whereas *Asc*^−/−^ mice are defective for IL-18 release but not pyroptotic cell death or IEC expulsion (Rauch et al., 2017) (Figures 1C, 1E, 2B). We also crossed OvaFla Villin-Cre-ER^T2^ mice to pyroptosis-deficient *Gsdmd*^−/−^ mice and found that they were defective for IL-18 release *in vivo* (Figures 1C, 1E).

Because *Nlrc4*^−/−^ IECs fail to undergo pyroptosis or IEC expulsion (Rauch et al., 2017), we noted that cells expressing the OvaFla transgene accumulate to much higher levels in the *Nlrc4*^−/−^ mice than in WT, *Asc*^−/−^, or *Gsdmd*^−/−^ mice, in which IEC expulsion still occurs (Figure 2). Higher levels of Ova antigen in IECs has previously found to correlate with higher levels of OT-I expansion in the spleen and mesenteric lymph nodes of mice (Vezys, Olson, & Lefrancois, 2000). Because of the differences in antigen burden, comparisons of *Nlrc4*^−/−^ mice to the other genotypes must be made with caution. Nevertheless, we found that OT-I T cells in the *Nlrc4*^−/−^ OvaFla mice divide and are activated at similar levels to the WT OvaFla mice following tamoxifen administration (Figure 3B–D). This activation occurred even when direct presentation of the OT-I peptide by IECs was eliminated on the K^bm1^ background (Figure 4B–C). These results are surprising for two reasons. First, it is not clear how IEC-derived antigens would be delivered to APCs in the absence of inflammasome-induced cell death. Other studies have suggested that IEC apoptosis, which may occur during homeostatic IEC turnover (Bullen et al., 2006; Hall, Coates, Ansari, & Hopwood, 1994; Marshman, Ottewell, Potten, & Watson, 2001; Shibahara et al., 1995; Watson et al., 2005), can be a source of antigen for T cell activation (Cummings et al., 2016; Huang et al., 2000). However, apoptotic IECs are expelled apically into the intestinal lumen (Bullen et al., 2006; Hall et al., 1994; Marshman et al., 2001; Shibahara et al., 1995; Watson et al., 2005), and so the exact mechanism of basolateral antigen delivery remains unclear—though it may involve luminal sampling by intestinal phagocytes (Farache et al., 2013) and/or the transfer of plasma membrane components (trogocytosis) (Dance, 2019). Cummings *et al* suggested that IECs can be engulfed by APCs, resulting in antigen presentation on MHC class II to induce CD4^+^ T regulatory cells, but this work did not examine antigen-specific responses or MHC class I presentation to CD8^+^ T cells. Further work is therefore needed to understand mechanisms of IEC-derived antigen presentation in the absence of inflammatory cell death. The second reason we were surprised to see CD8^+^ T cell activation in *Nlrc4*^−/−^ OvaFla mice is that these mice are presumably unable to produce inflammatory signals necessary to induce APC activation. However, previous studies have shown that OT-I T cells can be activated from constitutively expressed Ova in the absence of inflammation. In this scenario, the CD8^+^ T cells go on to become anergic and are likely eventually deleted from the periphery (Kurts et al., 1996; Kurts et al., 1997; W. Liu, Evanoff, Chen, & Luo, 2007; Vezys et al., 2000).

Since WT, *Asc*^−/−^, and *Gsdmd*^−/−^ IECs all undergo cell death and IEC expulsion in response to NLRC4 activation, these mice exhibit similar levels of OvaFla transgene expression in IECs, allowing for comparisons between these mouse strains. We found that OvaFla production leads to CD8^+^ T cell expansion and activation in all these strains. The expansion is at least partially dependent on gasdermin D, as *Gsdmd*^−/−^ OvaFla mice have significantly fewer OT-I T cells than their WT counterparts (Figure 3B, C). Interestingly, ASC-deficient OvaFla mice— in which IECs still undergo pyroptosis following NAIP–NLRC4 activation (Rauch et al., 2016)—show similar, or even increased, OT-I numbers in their tissues relative to WT OvaFla mice. These data, combined with the fact that *Gsdmd*^−/−^ and *Asc*^−/−^ OvaFla mice have little to no detectable IL-18 in their serum (Figures 1C, 1E), suggest that the difference in OT-I T cell proliferation between these strains is in some way related to pyroptotic antigen release. One hypothesis is that the gasdermin D pore, which has been shown to provide a lysis-independent portal for IL-1β, IL-18, and other small proteins (DiPeso et al., 2017; Evavold et al., 2018; Heilig et al., 2018), may act as a channel for small antigens to escape IECs prior to cell expulsion.

Because *Gsdmd*-deficiency only modestly affected OT-I responses, our data additionally suggest that there may both GSDMD-dependent and GSDMD-independent pathways by which IEC antigens can be cross presented to CD8^+^ T cells. Because *Batf3^−/−^*-dependent cDC1s have a known role in cross-presenting IEC-derived antigen (Cerovic et al., 2015), we sought to determine if the cross presentation occurring in the OvaFla mice similarly relied on these cells. We compared bm1^+^ OvaFla mice that received B6 CD45.1 bone marrow with those that received bone marrow from *Batf3*-deficient mice (Figure 5A). To our surprise, we found OT-I T cells were cross-primed in the bm1^+^ WT OvaFla mice, even in the recipients that lacked cDC1s (Figure 5A–D). Interestingly, these data contrast with the bm1^+^ *Nlrc4*^−/−^ OvaFla mice, where the recipients given *Batf3*-deficient bone marrow had significantly fewer activated OT-I T cells than their counterparts given *Batf3*-sufficient bone marrow. OT-I T cell activation in the bm1^+^ *Gsdmd*^−/−^ OvaFla mice partially relied on *Batf3*^+^ DCs. Furthermore, CellTrace Violet data show the OT-I T cells in the bm1^+^ *Nlrc4*^−/−^ and bm1^+^ *Gsdmd*^−/−^ OvaFla mice undergo fewer rounds of division in the absence of *Batf3* cDC1s (Figure 5D). These data suggest there may be two possible cross presentation pathways for IEC-derived antigen: one that occurs in the presence, and one in the absence, of inflammatory signals. Indeed, recent work shows that cDC2s can take on characteristics of cDC1s under inflammatory conditions (Bosteels et al., 2020) or in the absence of *Batf3* (Lukowski et al., 2021), though it remains uncertain if these cells are able to cross prime CD8^+^ T cells or provide T cells with the appropriate homing signals. Perhaps IL-18 and/or other inflammatory signals downstream of NAIP–NLRC4 activation can activate cDC1-like capabilities of cDC2s in our OvaFla system.

Our work raises several interesting questions for future study, including the mechanism of cDC maturation. The traditional model of DC maturation involves TLR signaling on the DC (Dalod, Chelbi, Malissen, & Lawrence, 2014). IL-1R (Pang, Ichinohe, & Iwasaki, 2013) or IL-18R (Li et al., 2004) on these cells might also trigger maturation, though further investigation is needed to understand how IL-1β, IL-18, or other inflammatory signals, such as eicosanoids (McDougal et al., 2021; Rauch et al., 2017), downstream of inflammasome activation might drive maturation of DCs that have acquired IEC-derived antigen.

Overall, our studies show that show that IEC-derived antigens are cross-presented both following NAIP–NLRC4 activation and under apparent homeostatic conditions (in the absence of NAIP–NLRC4 induced inflammation). In the context of NAIP–NLRC4 activation, cross-priming of CD8^+^ T cells is partially dependent on gasdermin D-mediated pyroptosis and can occur in the absence of cDC1s. These data add insights to the complex interactions between innate and adaptive immune responses occurring in the intestine.

## Materials and Methods

### Animals

All mice were maintained under specific pathogen-free conditions and, unless otherwise indicated, fed a standard chow diet (Harlan irradiated laboratory animal diet) *ad libitum*. OvaFla mice were generated as previously described (Nichols et al., 2017) and crossed to Villin-Cre-ER^T2^ (el Marjou et al., 2004) (Jax strain 020282). OvaFla Villin-Cre-ER^T2^ mice were additionally bred to *Gsdmd^−/−^*, *Asc*^−/−^ and *Nlrc4*^−/−^ mice. *Nlrc4*^−/−^ and *Asc*^−/−^ mice were from V. Dixit (Mariathasan et al., 2004) (Genentech, South San Francisco, CA). *Gsdmd*^−/−^ mice were previously described (Rauch et al., 2017). OT-I *Rag2*^−/−^ mice (from E. Robey, Berkeley, CA) were used as a source of OT-Is for all adoptive transfer experiments.

For chimera experiments, the above OvaFla lines were crossed to B6.C-*H-2K^bm1^*/ByJ mice (Schulze et al., 1983) (Jax strain 001060). For the bone marrow donors, B6.CD45.1 (Jax strain 002014) and *Batf3*^−/−^ (Jax strain 013755), mice were used.

Mice used for non-chimera experiments were 8-12 weeks old upon tissue harvest, and mice used as chimeras were 16-20 weeks old upon tissue harvest. Female mice were cohoused, and all experimental mice were age- and sex-matched when possible. OvaFla-only and Cre-only controls were littermates of the experimental mice. All animal experiments and endpoints were approved by and performed in accordance with the regulations of the University of California Berkeley Institutional Animal Care and Use Committee.

### Adoptive transfer of OT-I T cells

The spleen and mesenteric lymph nodes were harvested from OT-I *Rag2*^−/−^ mice, mashed between the frosted ends of two glass slides to create a single cell suspension, filtered through 100mm nylon mesh, and pooled into a single tube. Red blood cells were lysed with ACK Lysing Buffer (Gibco; A10492-01). Cells were labeled with CellTrace Violet (ThermoFisher; C34557) following the manufacturers protocol and transferred i.v. to mice anesthetized with isoflurane at 2×10^4^ cells per mouse.

### Tamoxifen administration

The tamoxifen chow used in these studies was purchased from Envigo (https://www.envigo.com/tamoxifen-custom-diets; 120856). The diet contains 250 mg of tamoxifen per kilogram of chow and was irradiated prior to shipping. Mice were fed *ab libitum* for two days for the experiments in Figures 1B–1C and one day for the remaining experiments. Envigo assumes approximately 40 mg of tamoxifen is consumed per kilogram of body weight per day for each mouse, though feed aversion leads to variable and limited initial food intake (Chiang et al., 2010).

### Flow cytometry

Spleens and mesenteric lymph nodes were harvested from euthanized mice and stored on ice in T cell media: RMPI 1640 (Gibco; 21870092) containing 10% FBS (Gibco, Cat#16140071, Lot#1447825), 1% penicillin-streptomycin, 1% L-glutamine, 1% sodium pyruvate, 0.5% 2-mercaptoethanol, and 25mM HEPES. For lymphocyte staining, tissues were mashed between the frosted ends of glass slides and filtered through 100mm nylon mesh. For myeloid staining, tissues were minced with scissors and forceps and incubated in T cell media containing 1 mg/mL collagenase VIII (Sigma; C2139-1G) at 37 °C for 45 minutes. The digested tissues were then passed through 70 mm filters and washed with T cell media. For all stains, red blood cells were lysed from a single cell suspension using ACK Lysing Buffer. Cells were counted using a Beckman Vi-CELL XR Cell Viability Analyzer (Beckman Coulter, Brea, CA), and 3×10^6^ cells per tissue per mouse were added to individual FACS tubes or wells of a 96-well non tissue culture treated round bottom plate.

For extracellular surface staining, cells were blocked for 20-30 minutes with a 1:1000 dilution of anti-mouse CD16 and CD32 antibodies (eBioscience; 14-0161-85) at 4 °C and then stained with a cocktail of antibodies for extracellular markers (Supplemental Table 1) at RT for 1 hour. All dilutions and washes were done with 1X PBS (Gibco; 10010049) containing 5% FBS/FCS.

For intracellular cytokine analysis, cells were incubated at 1×10^6^ cells/mL T cell media plus 1 μg/mL phorbol myristate acetate (PMA) (Invivogen; tlrl-pma), 1μg/mL ionomycin (Calbiochem; 407952-1MG), and 1 μg/mL GolgiPlug™ (BD Biosciences; 555029) at 37°C for 5 hours. Cells were then washed and blocked for 20-30 minutes with a 1:1000 dilution of anti-mouse CD16 and CD32 antibodies at 4 °C, and a surface stain was applied for 1 hour at RT (Supplemental Table 1). Cells were then fixed in 100mL eBioscience™ IC Fixation Buffer (Thermo; 00-8222-49) for 20-60 minutes RT, and then stained with an intracellular staining cocktail (Supplemental Table 1) in 1X eBioscience™ Permeabilization Buffer (Thermo; 00-8333-56) at RT for 1 hour. Cells were washed and resuspended in PBS prior to analysis. The data were collected on a BD Biosciences Fortessa (San Jose, CA) in the UC Berkeley Cancer Research Laboratory Flow Cytometry facility, and analysis was performed using FlowJo 10 Software (BD Biosciences, San Jose, CA).

### Generation of bone marrow chimeras

Eight- to twelve-week-old mice were lethally irradiated with a Precision X-Rad320 X-ray irradiator (North Branford, CT) using a split dose of 500 rads and then 450 rads, approximately 15 hours apart. Bone marrow was harvested from the long bones of the indicated donor strains, red blood cells were lysed using ACK Lysing Buffer, and CD3^+^ cells were depleted from the donor cells using a biotinylated anti-mouse CD3e mAb (BioLegend; 100304) and the Miltenyi MACS® MicroBead (Miltenyi; 130-105-637) magnetic depletion protocol with LD columns (Miltenyi; 130-042-901) to reduce graft vs host reactions (Selvaggi, Ricordi, Podack, & Inverardi, 1996). Recipient mice were anesthetized with isoflurane, and approximately 5×10^6^ donor cells were injected retro-orbitally. Females from the different strains were co-housed, and at least eight weeks passed between reconstitution and the start of any experiment.

### Immunofluorescence

Mice were fed a single day pulse of tamoxifen chow and euthanized two days from start of the chow feeding. Approximately 2.5 cm pieces were taken from the proximal and distal ends of the small intestine. These pieces were flushed and fixed in PLP buffer (0.05 M phosphate buffer containing 0.1 M L-lysine [pH 7.4], 2 mg/mL NaIO_4_, and 1% PFA) overnight at 4 °C. The following day, tissues were washed 2x in phosphate buffer and placed in 30% sucrose overnight at 4 °C. Tissue was frozen in Tissue-Tek® OCT (VWR; 25608-930), cut on a Leica cryostat, and sections were placed on Fisherbrand™ Tissue Path Superfrost™ Plus Gold Slides (Fisher Scientific; 15-188-48).

For staining, slides were allowed to warm to room temperature, traced with an ImmEdge Hydrophobic Barrier Pen (Vector Labs; H-4000), washed 3× in 1× PBS with 0.5% Tween-20, and blocked with 10% normal donkey serum (Sigma; D9663) in 0.5% Tween-20, 100 mM TrisHCl [pH 7.5], 150 mM NaCl, 0.5% blocking reagent (Perkin Elmer; FP1020) for 30 minutes. Tissues were then stained with 1:300 GFP polyclonal antibody (Invitrogen; A-6455) overnight at 4 °C. Slides were washed 3X and stained with donkey anti-rabbit Alexa Fluor 488 (Jackson Immunoresearch; 711-545-152) for 60 minutes at RT, followed by 150 nM Acti-stain™ 555 phalloidin (Cytoskeleton, Inc; PHDH1-A) and 100 mM DAPI (D1306) for 30 minutes at RT. Slides were then washed 2X in H_2_0 and sealed under glass coverslips prior to imaging. All antibody dilutions were done in 100 mM TrisHCl [pH 7.5], 150 mM NaCl, 0.5% blocking reagent; all washes were done in 1X PBS with 0.5% Tween-20.

Slides were imaged on a Zeiss LSM710 at the CNR Biological Imaging Facility at the University of California, Berkeley. Images were blinded and manually quantified for GFP^+^ IECs. For quantification, DAPI^+^ IECs were counted in at least 15 villi per mouse—DAPI^+^ cells were counted prior to revealing the GFP^+^ cells in the 488 channel. ImageJ (National Institutes of Health) was used to visualize images and globally adjust contrast and brightness for print quality following quantification.

### Serum IL-18 measurement

Thermo Scientific Immuno MaxiSorp ELISA plates (Thermo Fisher; 439454) were coated with 1μg/mL anti-mouse IL-18 mAb (MBL; D048-6) overnight at 4°C, and blocked with 1× PBS containing 1% BSA for 2-4 hours at RT. Serum was diluted 1:5 in PBS with 1% BSA, added to the plate with a purified IL-18 standard, and incubated overnight at 4°C. A biotinylated anti-mouse IL-18 sandwich mAb (BioXcell; BE0237) was added at 1:2000 in PBS with 1% BSA and incubated for 1-2 hours at RT. BD Pharmingen™ Streptavidin HRP (BD Biosciences; 554066) was added at 1:1000 in PBS with 1% BSA. Following a final 5× wash, plates were developed with 1 mg/mL OPD (Sigma; P3804-100TAB) in citrate buffer (PBS with 0.05 M NaH2PO4 and 0.02 M Citric acid) plus 9.8M H_2_O_2_. The reaction was stopped with a 3 M HCl acid stop after approximately 10 minutes. Absorbance at 490 nm was measured on a Tecan Spark® multimode microplate reader (Tecan Trading AG, Switzerland).

### Statistical analysis

For all bar graphs, data are shown as mean ± SD. For Figures 1–4, the significance between each genotype was calculated using one-way ANOVA and Tukey’s multiple comparisons test. For Figure 5, the significance between mice that received B6 CD45.1 and mice that received *Batf3*^−/−^ bone marrow was calculated using one-way ANOVA and Šídák’s multiple comparisons test. For all data, \**p* < 0.05, \*\**p*<0.01, \*\*\**p*<0.001. Tests were run using GraphPad Prism (San Diego, CA).

## Acknowledgements

We thank members of the Vance and Barton Labs for discussions, Greg Barton and Ellen Robey for comments on the manuscript, Dmitri Kotov for comments on the manuscript and advice, and Roberto Chavez for technical assistance. We also thank the UC Berkeley Cancer Research Laboratory Flow Cytometry facility, including Hector Nolla and Alma Valeros. Figure 4A was created with BioRender.com. Research reported in this publication was supported in part by the National Institutes of Health S10 program under award number 1S10RR026866-01. The content is solely the responsibility of the authors and does not necessarily represent the official views of the National Institites of Health. R.E.V. is an Investigator of the Howard Hughes Medical Institute, and research in his laboratory is funded by NIH grants AI075039, AI063302, and AI155634.

---

**Supplementary table 1.**
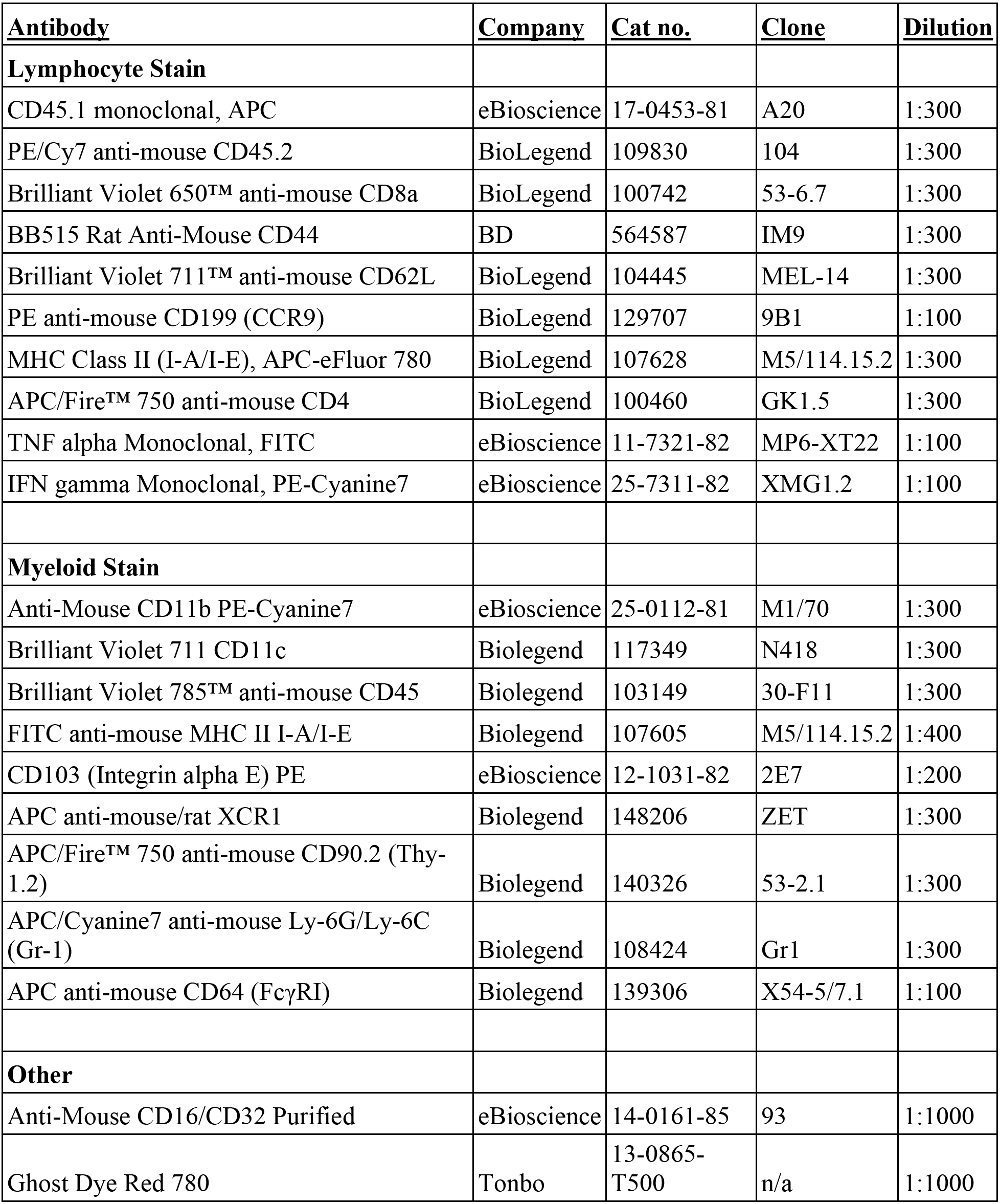
List of antibodies used for flow cytometry.

**Figure 3–figure supplement 1.**
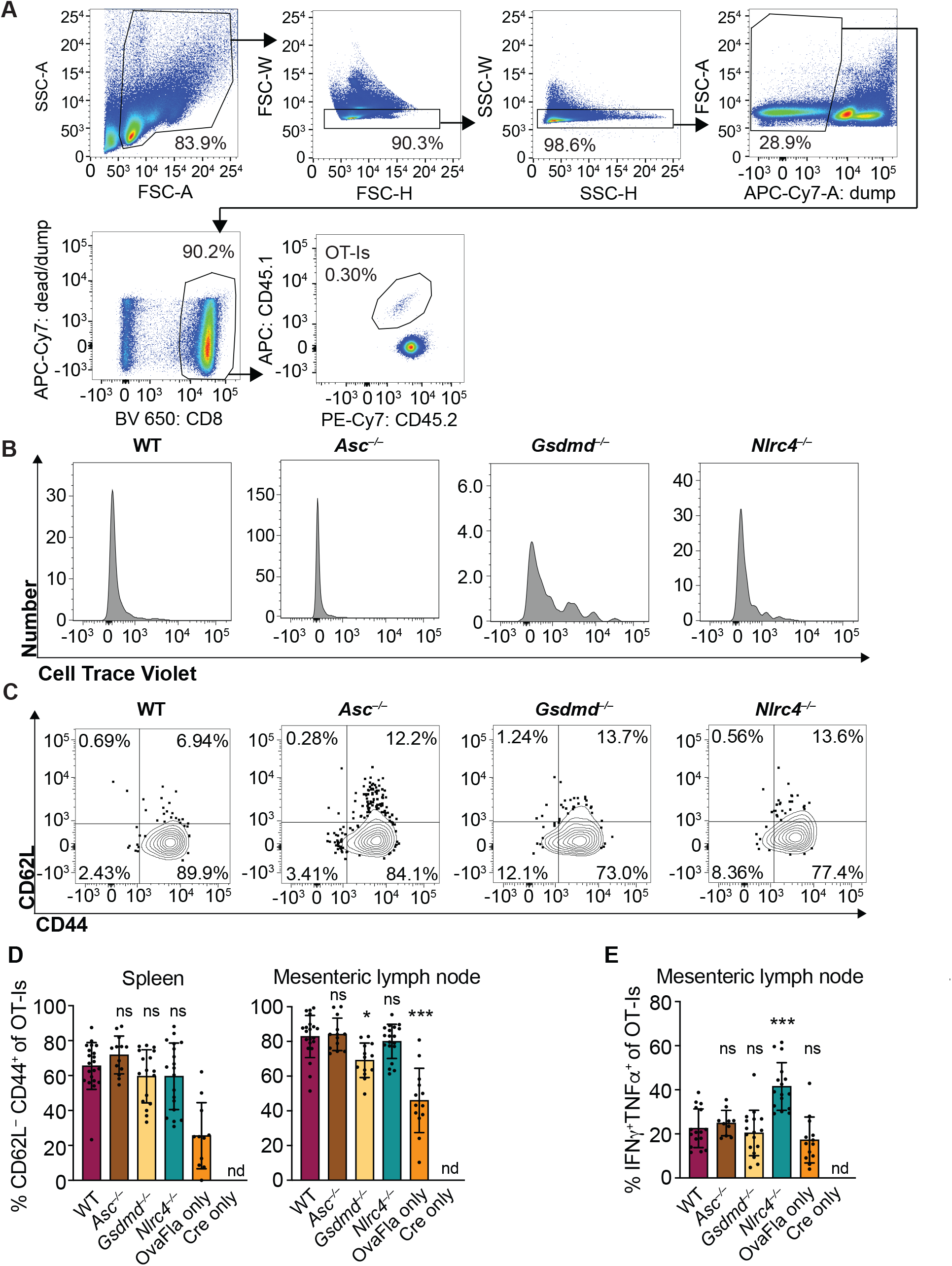
OvaFla expression in IECs results in OT-I proliferation and activation that is independent of ASC and NLRC4 but partially dependent on gasdermin D. **A.** Flow cytometry gating strategy for identifying OT-I T cells. **B.** Representative histograms of CellTrace Violet dilution for each of the OvaFla mouse lines. **C.** Representative dot plots of each OvaFla mouse line showing the gating strategy for identifying CD62L–CD44^+^ OT-Is. **D**. Percent of OT-Is that are CD64L^−^CD44^+^ per spleen (left) and mesenteric lymph node (right). **E**. Percent of OT-Is from the mesenteric lymph node that are IFN γ^+^TNFa^+^ following a 5-hour ex vivo stimulation with PMA (1 μg/mL) and ionomycin (1 μg/mL). D-E, data are from three independent experiments, and each dot represents an individual mouse. Data shown as mean +/– SD. Significance calculated using one-way ANOVA and Tukey’s multiple comparisons test (\**p* < 0.05, \*\**p* < 0.01, \*\*\**p* < 0.001). Only *p* values between WT and other experimental groups are shown.

**Figure 4–figure supplement 1.**
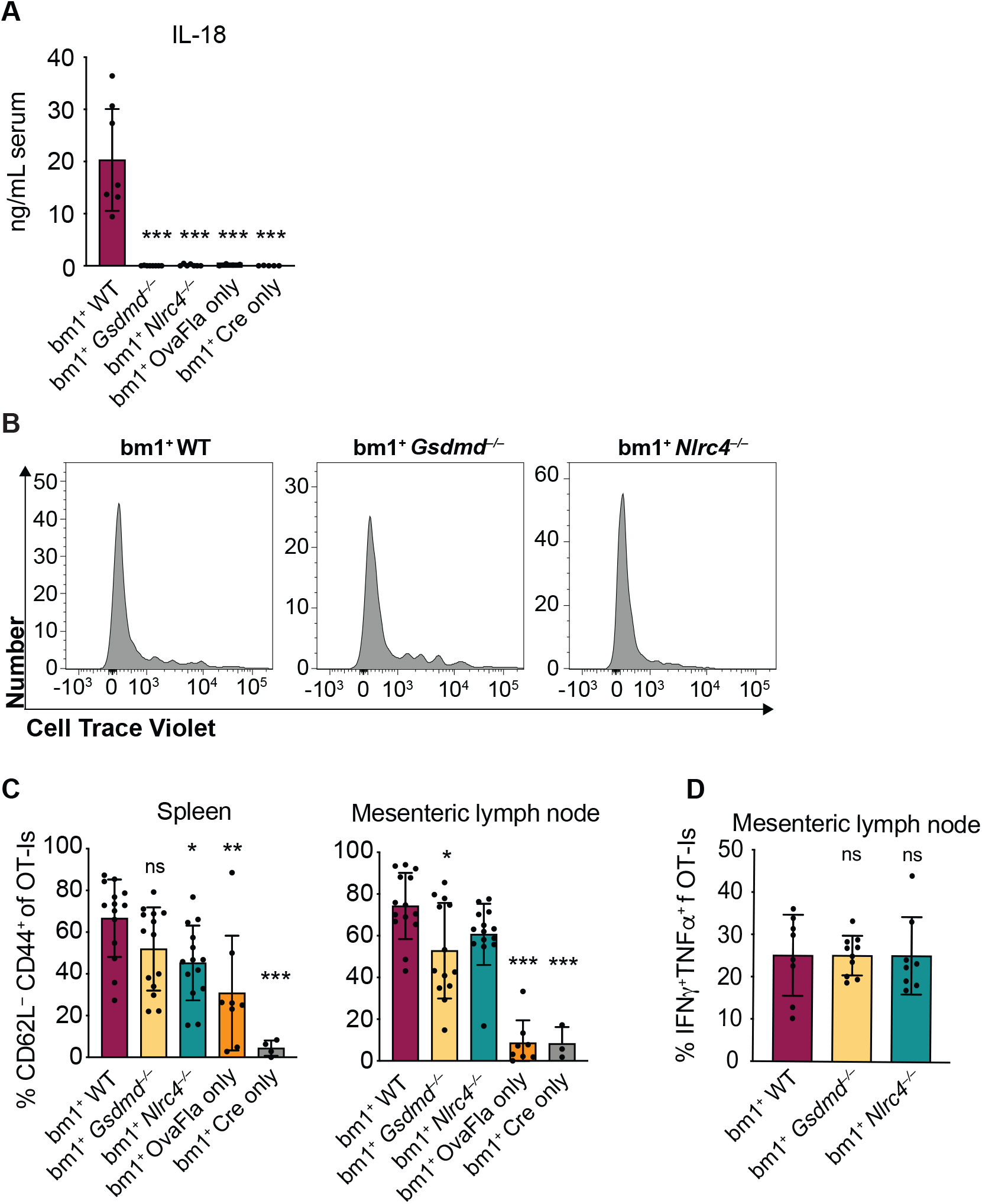
OvaFla expression in IECs of bm1^+^ OvaFla mice results in NAIP–NLRC4 expression and OT-I cross-priming that is independent of NLRC4 but partially dependent on gasdermin D **A.** Quantification of IL-18 ELISA performed on serum from the mice shown in Figure 4 at day 5 post tamoxifen chow start. **B.** Representative histograms of CellTrace Violet dilution for the indicated OvaFla mouse lines. **C.** Percent of OT-Is that are CD62L–CD44^+^ in the spleen (left) and mesenteric lymph nodes (right) of the mice shown in Figure 4. **D.** Percent of OT-Is from the mesenteric lymph nodes of the mice in Figure 4 that are IFNγ^+^TNFα^+^ following a 5-hour ex vivo stimulation with PMA (1 μg/mL) and ionomycin (1 μg/mL). A, data are from two independent experiments. C, data are from three independent experiments. Each dot represents an individual mouse. Data shown as mean +/– SD. Significance calculated using one-way ANOVA and Tukey’s multiple comparisons test (\**p* < 0.05, \*\**p* < 0.01, \*\*\**p* < 0.001). Only *p* values between WT and other experimental groups are shown.

**Figure 4–figure supplement 2.**
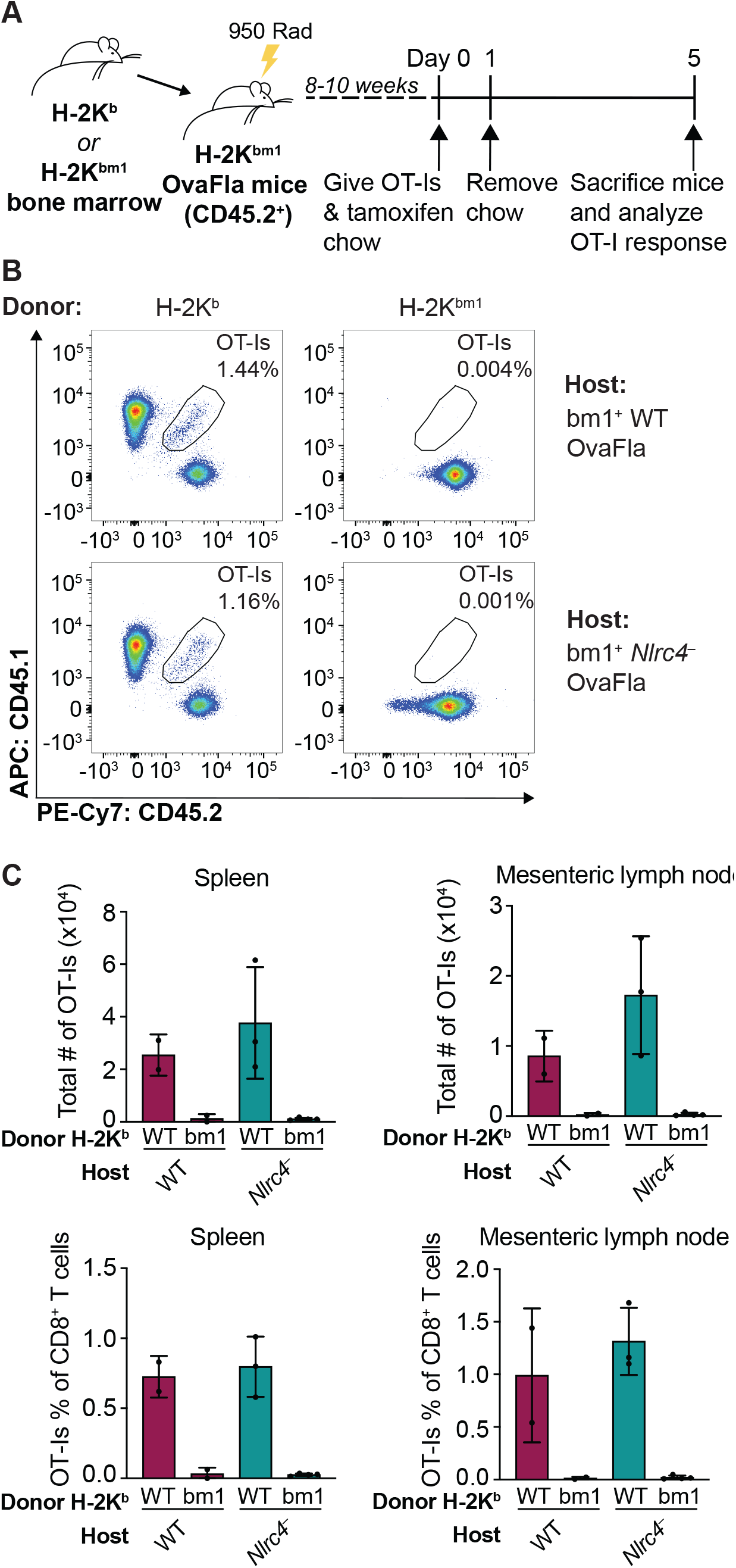
K^b^ donor bone marrow is required for OT-I proliferation and activation in bm1^+^ OvaFla bone marrow chimeras. **A.** Schematic depicting the production and analysis workflow of chimeric bm1^+^ OvaFla mice that were given either B6 H-2K^b^ or H-2K^bm1^ bone marrow. **B.** Representative flow plots demonstrating the absence of OT-Is in the mice given H-2K^bm1^ bone marrow, as depicted in A. **C.** Quantification of the total number of OT-Is (top) and the OT-Is as a percent of total CD8+ T cells (bottom) in the spleen (left) and mesenteric lymph nodes (right) of bm1^+^ WT and bm1^+^ Nlrc4^−^ OvaFla mice as depicted in A. Data are from a single experiment, and each dot represents an individual mouse. Data shown in C as mean +/– SD.

**Figure 5–figure supplement 1.**
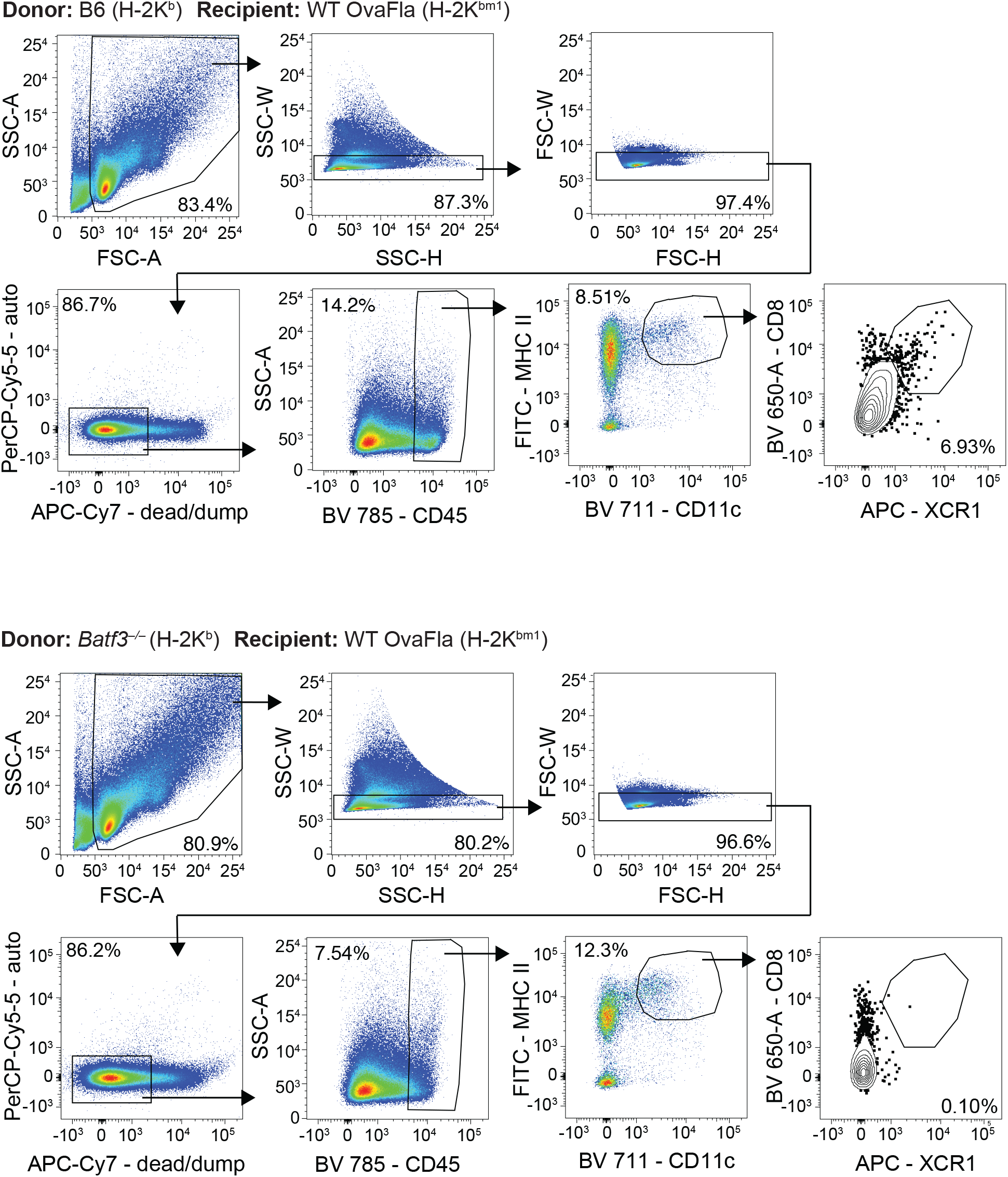
Gating demonstration for Figure 5A. Representative dot plots from one bm1^+^ WT OvaFla mouse that received B6 bone marrow (top) and one bm1^+^ WT OvaFla mouse that received *batf3*^−/−^ bone marrow (bottom).

**Figure 5–figure supplement 2.**
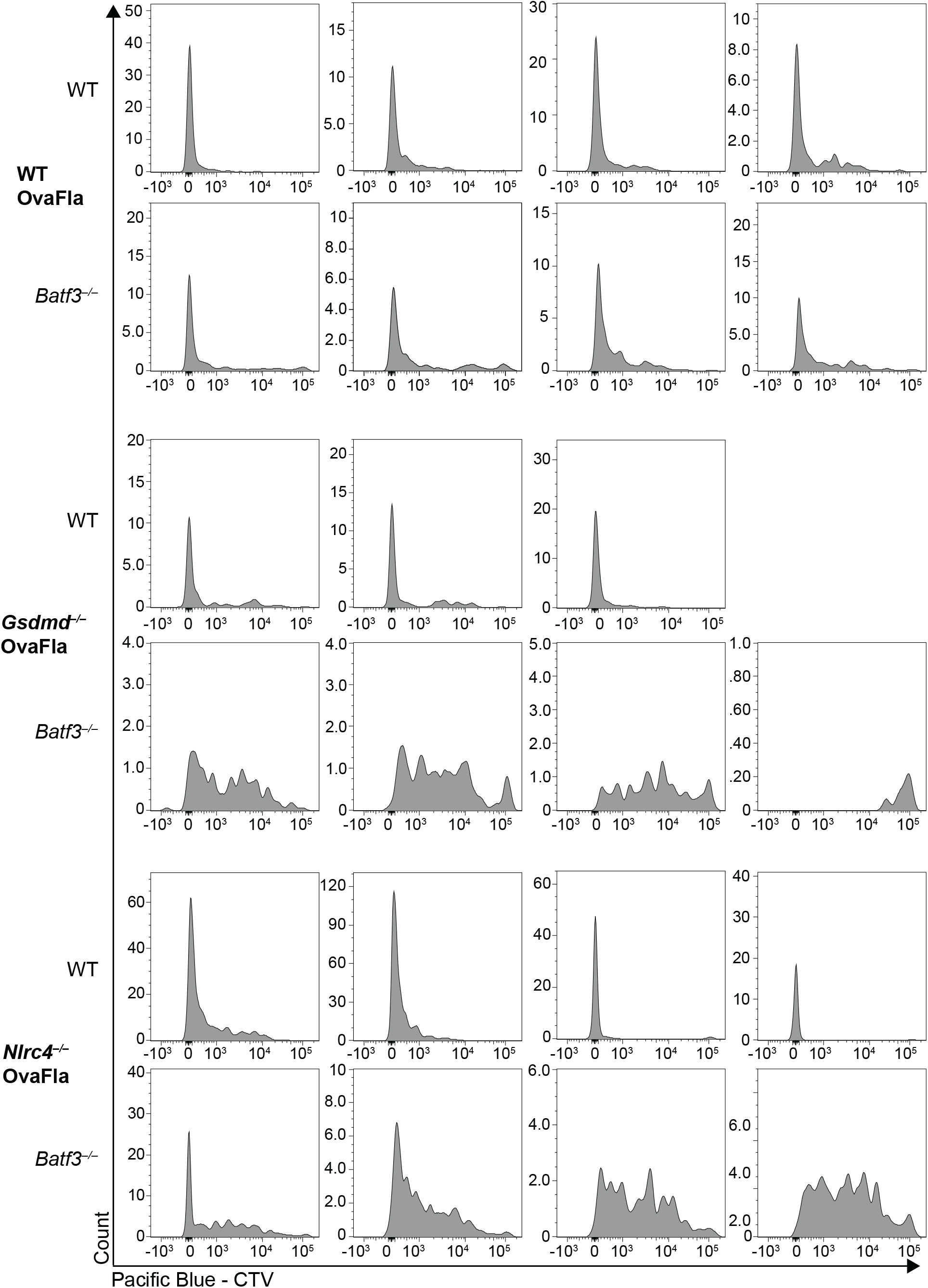
Representative histograms demonstrating the dilution of CellTrace Violet dye of OT-Is from individual mice shown in figure 5D. OT-Is are gated per Figure 3–figure supplement 1.

**Figure 5–figure supplement 3.**
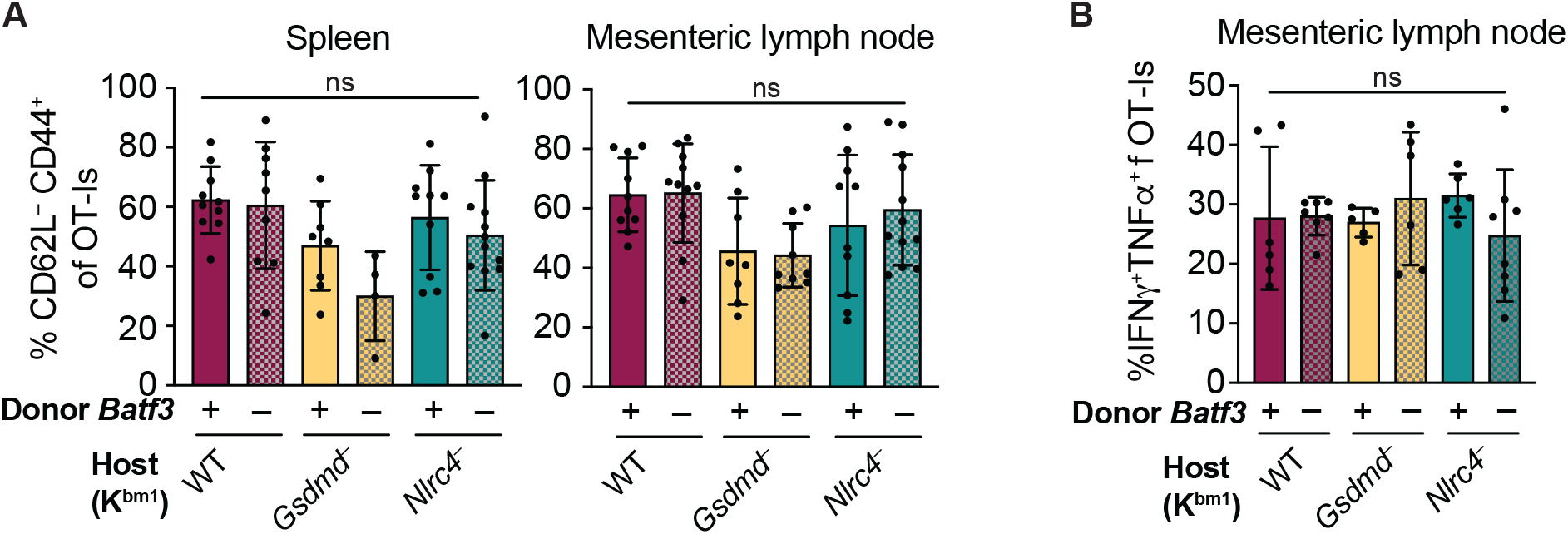
No difference in the percent of CD62L^−^CD44^+^ OT-I T cells or in the IFNγ and TNFα production between genotypes of bm1^+^ OvaFla mice. **A.** Percent of OT-Is that are CD62L^−^ CD4^4+^ in the spleen (left) and mesenteric lymph nodes (right). **B.** Percent of OT-Is from the mesenteric lymph node that are IFNγ^+^ and TNFα^+^ following a 5-hour ex vivo stimulation with PMA (1 μg/mL) and ionomycin (1 μg/mL). A, data are from three independent experiments. B, data are from two independent experiments. Each dot represents an individual mouse. Data shown as mean +/– SD. Significance calculated using one-way ANOVA and Šídák’s multiple comparisons test (*p < 0.05, **p < 0.01, ***p < 0.001).

